# Neutrophil extracellular trap formation correlates with improved overall survival in ovarian cancer

**DOI:** 10.1101/862292

**Authors:** Besnik Muqaku, Dietmar Pils, Johanna C. Mader, Stefanie Aust, Andreas Mangold, Liridon Muqaku, Astrid Slany, Giorgia Del Favero, Christopher Gerner

## Abstract

It is still a question of debate whether neutrophils, often found in the tumor microenvironment, mediate tumor-promoting or rather tumor-inhibiting activities. The present study focusses on the involvement of neutrophils in high grade serous ovarian cancer (HGSOC). Multi-omics data comprising proteomics, eicosadomics, metabolomics, Luminex-based cytokinomics, and FACS data were generated from ascites samples. Integrated data analysis demonstrates a significant increase of neutrophil extracellular trap-(NET) associated molecules in non-miliary ascites samples. A co-association network analysis performed with the ascites data further revealed a striking co-correlation between NETosis-associated metabolites with several eicosanoids. Investigating primary neutrophils from healthy domors, NET formation was induced using ionomycin or phorbol ester. Data congruence with ascites analyses indicated the predominance of NOX-independent NETosis. NETosis is associated with S100A8/A9 release. An increase of the S100A8/CRP abundance ratio was found to correlate with improved survival of HGSOC patients. The analysis of additional five independent proteome studies with regard to S100A8/CRP ratios confirmed this observation. In conclusion, here we present evidence that increased NET formation relates to improved outcomes in cancer patients.

**Graphical abstract:** NETs releasing neutrophils through detaching of small tumor nods dictate the building of bigger in size and fewer in number of tumors in the non-miliary spreading tumor. Increased angiogenesis associated with increased blood circulation may contribute to less suppressive effects on NETs formation in the non-miliary tumor type. Tumor origin, *i.e*. fallopian tube for the miliary or ovary for the non-miliary tumors, may influence the angiogenesis and therewith – through facilitating of neutrophils activation – (co)determine the type of tumor spread.

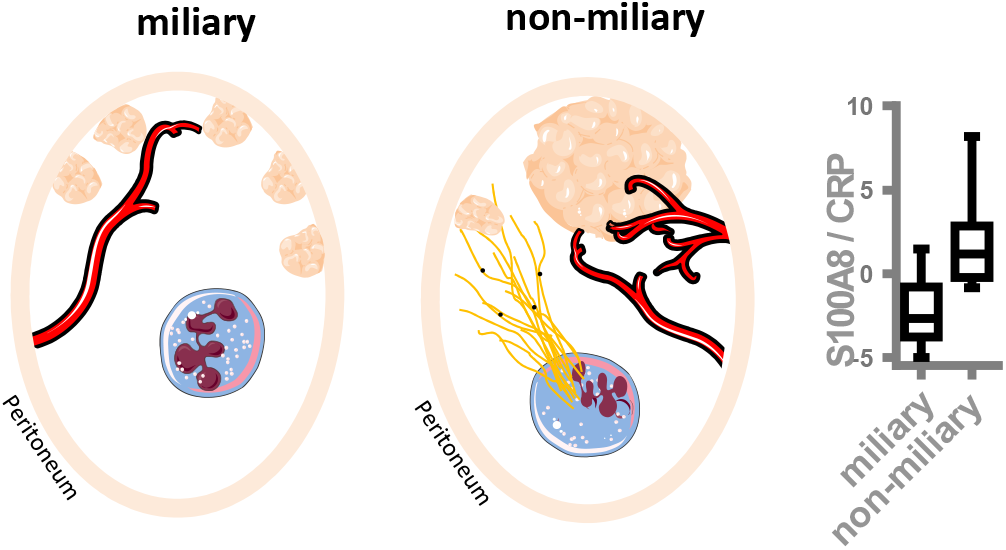

## Introduction

The tumor microenvironment, including the immune system, may have an important impact on tumor progression and treatment response [1, 2]. Focussing on tumor-stroma interactions, we are applying molecular profiling via mass spectrometry (MS), especially by the combination of proteomics, lipidomics and metabolomics [3–6]. This approach supports the analysis of pathomechanisms and related adaptative responses such as inflammation in great detail [7, 8]. With regard to high grade serous ovarian cancer (HGSOC), it allowed us to identify predictive marker profiles [9] as well as novel drug targets [10]. A distinct role of the immune system in ovarian cancer affecting the kind of tumor spread of ovarian cancer was described by us already previously [11].

High grade serous ovarian cancer still represents a major clinical challenge, the overall survival has only slightly improved within the last decades [12]. Peritoneal tumor spread and a unique tumor microenvironment provided by accumulated peritoneal fluid (ascites) characterize this disease. A classification based on macroscopic features distinguishes two types of peritoneal tumor spread in HGSOC [13]. The miliary type metastases appear as widespread and millet sized lesions, whereas non-miliary metastases are bigger in size, fewer in number and consist of exophytically growing implants. The first one is characterized by worst prognosis, increased systemic inflammation and little adaptive immune reactions in contrast to non-miliary, which is probably spread via circulation and is more infiltrated by cells of the adaptive immune system [11, 14].

Among the members of the innate immune system, neutrophils belong to the first responders to infections, injury or damage-associated molecular patterns (DAMPs) [15, 16] and may eventually get activated by hypoxic conditions of the tumor microenvironment, as characteristic for the peritoneum in HGSOC [17, 18]. Neutrophils exert their functions through phagocytosis, degranulation or release of neutrophil extracellular traps (NETs)[19], a physiologic way of cell death called NETosis [17, 18, 20]. NETs are DNA structures associated with proteins such as histones and others with antibacterial activity, including elastase (ENOA), myeloperoxidase (MPO), cathepsin G (CTG) and lactoferrin (LTF) [20, 21]. Under normal conditions, after successful clearance of an infection, a resolution of inflammation is initiatiated. However, malignant pathological conditions may be accompanied by chronic and apparently disorganized inflammatory response [17, 18, 22]. While immune therapy aiming at fostering immune responses against cancer cells has revolutionized cancer therapy [23], we have also observed that immune cells may have strong tumor-promoting capabilities [24]. Actually, apparently conflicting data are available regarding harmful [25–27] or desirable [28, 29] effects of neutrophils in ovarian cancer.

The present study focusses on the potential contribution of neutrophils to mechanisms of tumor spread and overall survival in HGSOC patients. Integrating MS-based multi-omics analysis to ascites samples collected from HGSOC patients, we demonstrate the involvement of neutrophil-derived effector molecules in the pathoegenesis of this disease and present a marker profile indicative for the neutrophil status in relation to systemic inflammation which seems to be predictive for overall survival also in case of other tumor entities.

## Results

### Evidence for activated neutrophils in ascites of non-miliary type of tumor spread

Ascites samples were obtained from 18 patients as described previously [9] and classified according to macroscopic features upon surgery into miliary (11 patients) and non-miliary (7 patients). Figure 1A indicates proteins caracteristic for NETs according to Brinkmann et al. [20] which were found up-regulated in non-miliary ascites samples (Table S1). This observation indicates functional activation of neutrophils in patients with non-miliary tumor spread. Antioxidant proteins such as glutathione S-transferase P (CSTP1), glutathione S-transferase omega-1 (GSTO1) and glutathione synthetase (GSS) were also found up-regulated in non-miliary samples (Fig. 1A). In addition, the calprotectin constituents S100A8 and S100A9 were found up-regulated in these samples. As the inflammation marker calprotectin is derived from activated phagocytes, it is linked with local inflammation [30, 31]. In contrast, the liver-derived systemic inflammation markers C-reactive protein (CRP) and serum amyloid A-1 protein (SAA1) were found up-regulated in the miliary samples (Fig. 1A).

**Figure 1:**
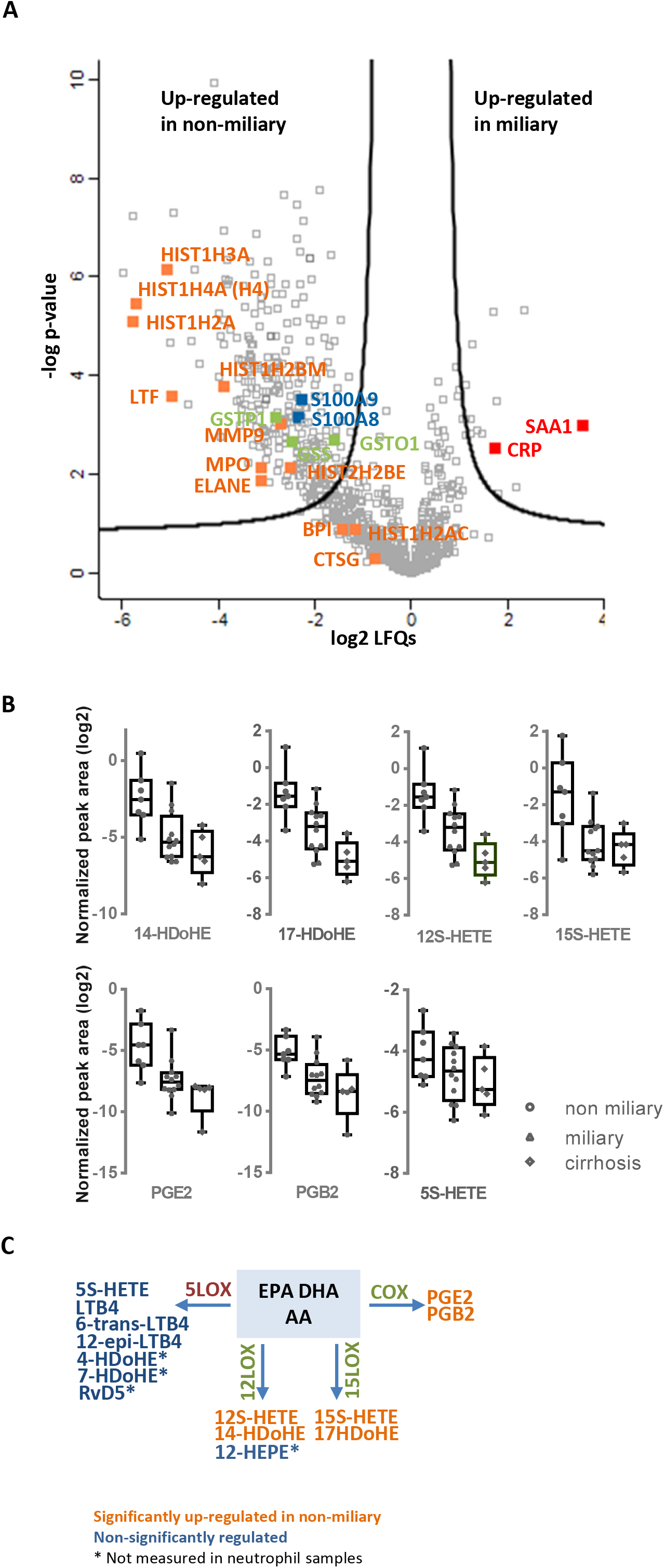
Untargeted multi-omics analysis of ascites samples. **A - Proteomics**, the volcano plot shows the results of comparative analysis between miliary and non-miliary ascites samples. Black lines represent threshold criteria for significant regulation with p-value < 0.05 and fold change > 2. NETs proteins are labeled in orange, whereas blue and red labeled proteins are related to local and systemic inflammation, respectively. Selected proteins linked to oxidative stress are labeled in green. **B - Eicosadomics**, boxplots show abundance levels at a logarithmic scale based on 2 of significantly regulated eicosanoids in non-miliary in comparison to miliary ascites samples (adjusted p-value < 0.05). In case of 5S-HETE, regulation was not significant. **C – Eicosanoid class switching**, three main precursors of eicosanoids are shown in the middle (AA, EPA, DHA) and eicosanoids are grouped dependent on enzymes from which they are synthesized (5-LOX, 12-LOX. 15-LOX and COX).

### Evidence for eicosanoids class switching in neutrophils from non-miliary samples

A comparative analysis of polyunsaturated fatty acids as well as their oxidation products performed by high-resolution mass spectrometry identified six eicosanoids (14-HDoHE, 17-HDoHE, 12S-HETE, 15S-HETE, PGE2 and PGB2) significantly up-regulated in the non-miliary compared to the miliary ascites samples (Fig. 1B and Table S2). Remarkably, these eicosanoids are products from 15-LOX, 12-LOX, and COX enzymes, whereas products of the 5-LOX enzyme [32] were positively identified, but not significantly regulated (Fig. 1B, 1C, S2 and Table 2S). As inflammatory stimulated neutrophils initially release mainly 5-LOX products, this finding suggested that neutrophils in the non-miliary samples had already switched to 12-LOX, 15-LOX and COX-products known to be involved in the resolution of inflammation [22] (Fig. 1C). The COX-product PGE2 (Fig. 1B) is actually known to further promote eicosanoid class switching in neutrophils initiating a feed-back loop [22, 33].

### Co-association network analysis revealed a strong correlation of six metabolites with eicosanoids

In order to learn more about the pathophysiological processes occurring in ovarian cancer patients, a co-association network analysis was generated. For that, the proteomics and eicosadomics data presented above were complemented with previously published data (metabolomics, transcriptomics, cyto/chemokine analyses and FACS data) [11]. The results show a molecular network signature with six metabolites (glutamate, aspartate, spermidine, spermine, taurine and histamine) establishing a hub in the center of the network predominantly correlating with eicosanoids. (Fig. 2). All six metabolites showed significantly increased concentrations in non-miliary compered to miliary ascites samples [11] (Fig. 2). Thus, a co-regulation of eicosanoids with metabolites was strongly suggested, motivating us to investigate the underlying pathomechanism. Remarkably, the ani-inflammatory cytokine IL-10 was found positively correlated with miliary samples.

**Figure 2:**
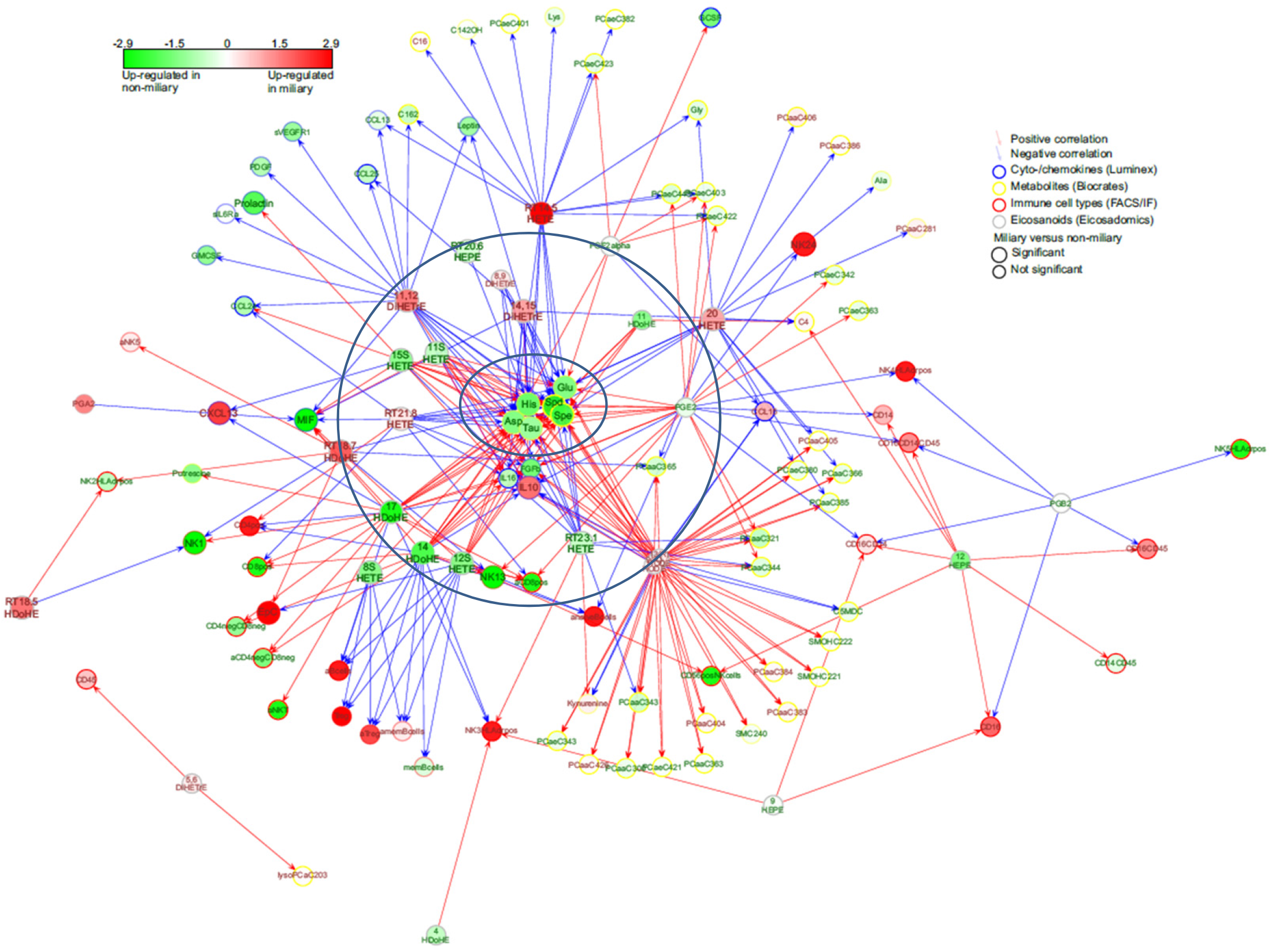
Network signature. A co-association network was built with molecules (eicosanoids, metabolites, proteins, RNA) and immune cells which were significantly regulated in non-miliary compared to miliary ascites samples. Six metabolites (Asp, Glu, taurine, histamine, spermidine, spermine) within inner circle built a hub in the middle of the network and were strongly regulated by many eicosanoids along outer circle.

### Activated neutrophils may account for most molecular alterations observed in ascites samples

To verify whether activated neutrophils might represent a plausible source of deregulated molecules in the non-miliary ascites samples, we performed *in vitro* stimulatory experiments with neutrophils isolated from healthy donors. Formation of NETs in a process termed NETosis eventually resulting from strong neutrophil activation [18]. Following previous studies, isolated neutrophils were treated either with phorbol 12-myristate 13-acetate (PMA), inducing NOX-dependent NETosis, or ionomycin, inducing NOX-independent NETosis [34–37]. Proteome profiling demonstrated up-regulation of some NETs proteins indicated in Figure 1 already after one hour treatment, and upregulation of most indicated NETs proteins three hours after treatment with PMA or ionomycin, respectively (Fig. 3A). The effect obtained with ionomycin was apparently stronger compared to PMA.

**Figure 3:**
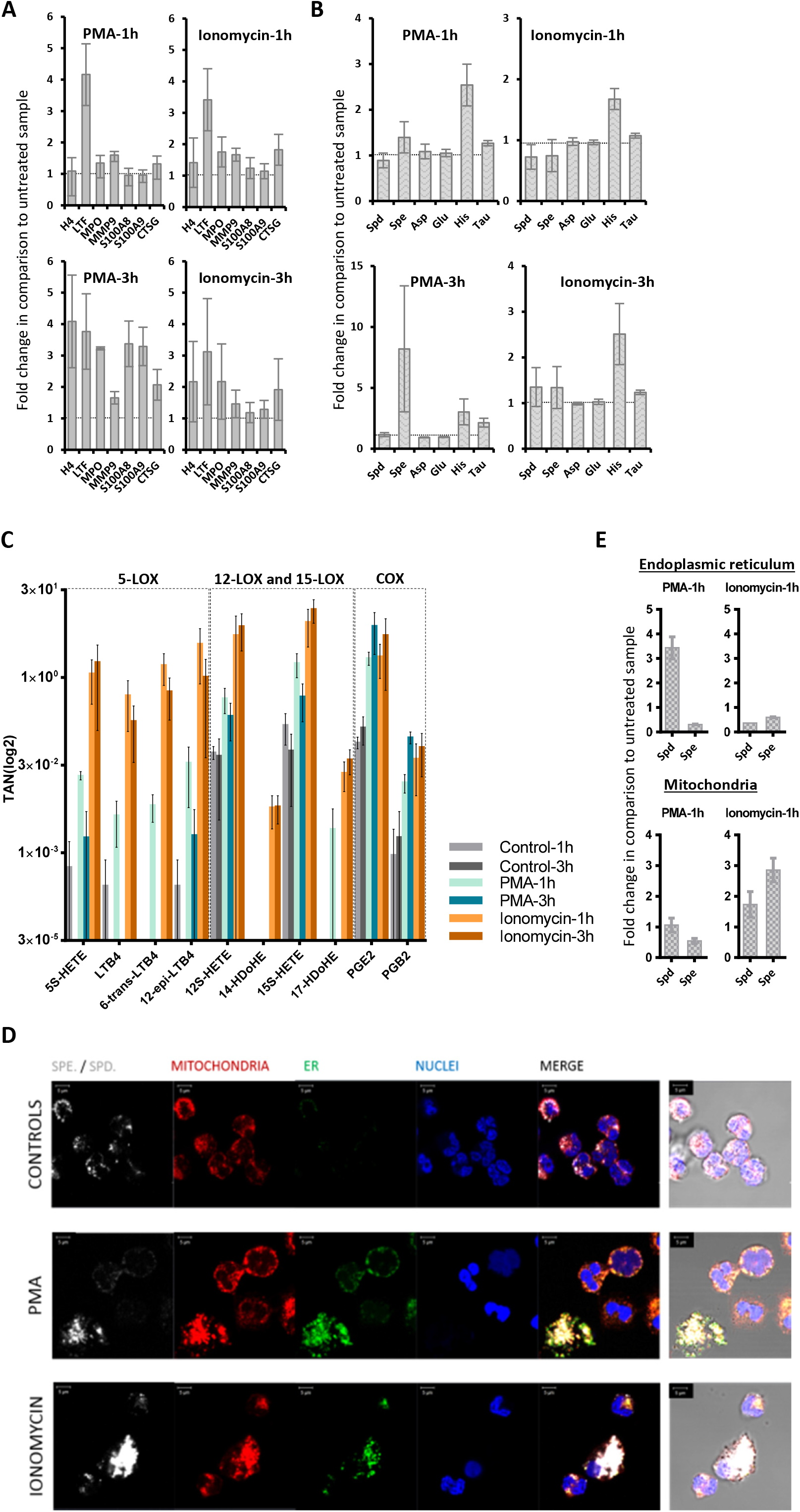
Targeted multi-omics analysis of neutrophils. Neutrophils isolated from healthy donors (n=3) were treated with PMA (25nM) or ionomycin (4μM). The cell supernatants were isolated and analyzed. Error bars indicate standard deviation. **A - Proteomics**, the fold change in comparison to untreated samples of NETs proteins and S1008/9 proteins is shown. **B - Metabolomics**, the regulation upon treatment of six metabolites as fold change compared to untreated samples is represented. A fold change value lower than 1 means down-regulation and a value higher than 1 indicates up-regulation. **C - Eicosadomics**, the abundance of selected eicosanoids as total area normalized to global deuterated standards (TAN) is blotted. Neutrophils (n=3) were cultured in RPMI medium supplemented with 0.01% FCS. **D - Immunolocalization of spermine/spermidine in neutrophils**. Comparison between untreated CONTROLS, or activation (PMA or IONOMYCIN, 1h). Spermine/spermidine (SPE./SPD.) is depicted in white, mitochondria (TOM20 protein) in red and endoplasmic reticulum (ER) in green. Nuclei are counterstained in blue with DAPI. Merged images are provided as fluorescence confocal images with or without transmitted light channel for visualization of cellular morphology in addition to the structures of interest. Scale bars stand for 5 μm. **E - Mitochondria and endoplasmic reticulum**, the organelles were isolated by sucrose gradient from lysed neutrophils. The fold change of spermidine (Spd) and spermine (Spe) in the endoplasmic reticulum (ER) as well as the mitochondria (Mit) by comparing treated and untreated cells are highlighted.

The metabolomics analysis of neutrophil supernatants, focusing on the six metabolites building a hub in the middle of the network, revealed up-regulation of spermine, spermidine, histamine, and taurine three hours after treatment (Fig. 3B). This suggests that neutrophil activation might also be related to the variations in the metabolic profile [11] (Fig. 2). Indeed, the positively charged polyamines have already been described to be released together with the NETs from activated neutrophils [38]. Furthermore, spermine has been described to attenuate mitochondrial swelling induced from high cytoplasmic calcium concentrations as caused by ionomycin treatment [39]. Thus, in order to verify if we could reproduce a similar response *in vitro*, we performed immunofluorescence experiments aimed to compare the subcellular localization of spermine/spermidine in neutrophils before and after activation. Apparently the activation with PMA induced the enlargement of ER, more powerful than ionomycin, while the polyamines were found to co-localize with both ER and mitochondria upon treatment (Fig. 3D). As anti-spermine antibodies cannot distinguish between spermine and spermindine, we extended this analysis with LC-MS analyses of sub-cellular fractions enriched in mitochondria and endoplasmic reticulum from untreated and stimulated neutrophils, respectively. Spermidine, but not spermine, was apparently strongly retained in the ER induced by PMA. Both polyamines were found upregulated in the mitochondrial fractions upon ionomycin treatment (Fig. 3E).

The biological functions of taurine, but also spermidine and spermine are related to antioxidant properties [40–42]. Upon inflammation-mediated oxidative stress, taurine may neutralize toxic oxidants generated from MPO by activated neutrophils [40]. Actually, neutrophils are known to contain very high concentrations of intracellular taurine [40]. Thus, the presently observed increased taurine concentrations may also be related to neutrophil activation resulting in NETosis in non-miliary ascites samples.

Also the eicosanoid analyses results of neutrophil supernatants are consistent with the suggested role of neutrophils in ascites samples. All six significantly regulated eicosanoids (Fig. S3A) were found induced upon stimulation with increasing values after one and three hours, respectively. While the 5-LOX products were found strongly induced upon one hour treatment, their abundance values was found consistently decreased after additional two hours (Fig. S3A). In contrast, the similarly induced 12/15-LOX and COX products did not decrease upon prolongued incubation. When using 0.01% FCS instead of 10% FCS, even more contrasting results were obtained due to high endogenous levels of eicosanoids detected in FCS (Fig. 3C). The results of the eicosadomics analysis were reproduced by repeating the experiment with ionomycin using neutrophils isolated from additional five healthy donors (Fig. S3B).

In summary, the *in vitro* experiments support the interpretation that activated neutrophils may represent the primary source of the observed alterations of eicosanoids and metabolites in ascites samples. Generally, the ionomycin-induced effects on neutrophils showed higher similarities to the molecular patterns observed in the ascites samples when compared to the observed effects using PMA, pointing to a rather NOX-independent activation pathway *in vivo*.

### Higher S100A8/CRP ratios in ascites samples positively correlate with overall survival

In clinical practice, elevated level of C-reactive protein (CRP) are often associated with unfavorable disease progression [43]. S100 proteins are regulated differently and thus show rather poor corrlations with CRP [44–46]. As CRP rather relates to systemic inflammation in contrast to S100 proteins, a ratio of these proteins may indicate to what extent inflammatory deregulation was becoming systemic and thus relate to overall survival.

Higher values of S100A8/CRP ratio were found in non-miliary samples described to have better prognosis (Fig. 4B). In order to compare the present data with a non-neoplastic but severe inflammatory disease, liver cirrhosis was included in the present considerations. Remarkably, the S100A8/CRP ratio was found at similar rates in patients with miliay tumor spread compared to patients with liver cirrhosis.

**Figure 4:**
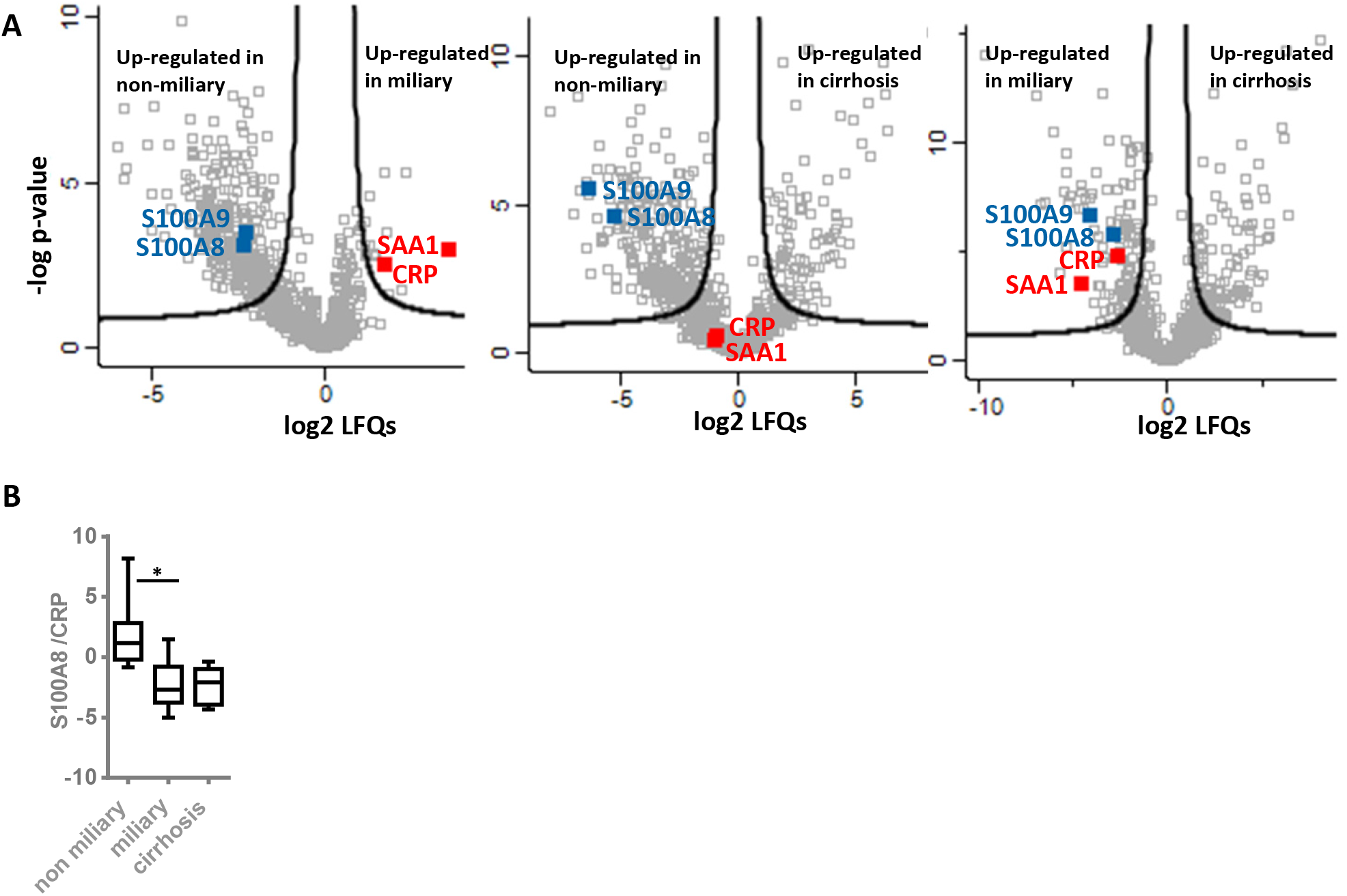
Inflammation marker expression and ratios. The volcano plots illustrate results of comparative analysis using shotgun proteomics data between three groups of ascites patients: non-miliary vs. miliary (left), non-miliary vs. cirrhosis (middle) and miliary vs. cirrhosis (right). Only selected proteins usually related to local-(blue) and systemic (red) inflammation are marked. Proteins above black lines are significantly regulated with FDR-value < 0.05 and fold change > 2 **(A)**. The abundance ratios within each patient group of data obtained from the shotgun analysis for S100A8 and CRP **(B)**.

### NETs proteins and S100A8/CRP ratio may serve as prognostic biomarker

To collect additional evidence for neutrophil activation in another kind of tumor and its potential association with overall survival, we re-evaluated published proteomics data from our laboratory generated from the analysis of cerebral melanoma metastases [47]. Melanoma patients (n=18), undergoing a MAPKi therapy, were classified depending on progression-free survival (PFS) in good (PFS ≥ 6 months) and poor responder (PFS ≤ 3 months). Comparing these two groups, the present analysis revealed significant up-regulation of several neutrophil-specific proteins, labeled in orange, in metastases isolated from good responders compared to those of poor responders (Fig.5A). Actually, similar to one hour neutrophils treatment (Fig. 3A), here we observed no release of histones but higher levels of CTSG suggesting neutrophil activation not necessarily resulting in NETosis. Anyhow, the molecular signatur of neutrophil activation was also associated with better prognosis in the melanoma patients.

**Figure 5:**
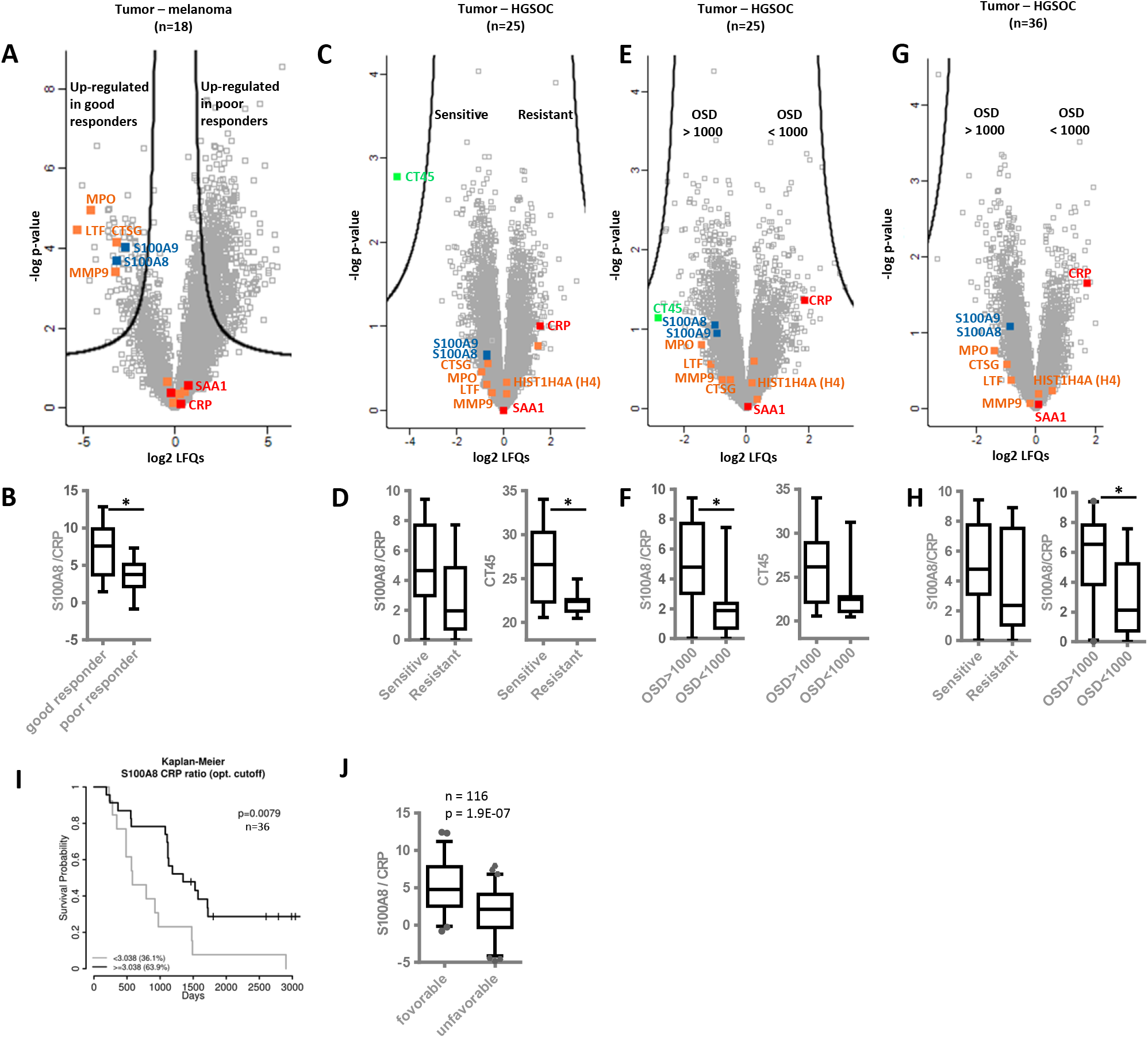
Shotgun proteomics analysis of tumor tissue samples isolated from melanoma and HGSOC patients. The volcano plot is showing the difference in protein abundances between cerebral metastases samples of melanoma patients, which upon MAPKi treatment were grouped in the poor (n=13, PFS ≤ 3 months) and good responder (n=5, PFS ≥ 6 months) **(A)**. The volcano plots represent the difference in protein abundances between omental tumor tissue samples of HGSOC patients grouped: dependent on chemotherapy treatment response in resistant (n=11) and sensitive (n=14) **(C)** and dependent on overall survival in the patients with 1000 < OSD > 1000 **(E, G)**. In all volcano plots, proteins above black lines were significantly regulated. The same significant regulation criteria were applied as reported in the original publications. Boxplots represent the S100A8/CRP abundance ratio in melanoma patient (B), HGSOC patient groped in resistant and sensitive **(D, H)** as well as HGSOC patients grouped in patients with 1000 > OSD < 1000 **(F, H)**. Additionally, the abundance distribution of CT45 protein (cancer/testis antigen 45) among four groups of HGSOC patients (n = 25) is shown **(D, F)**. Kaplan-Meier analysis of survival probability based on S100A8/CRP abundance ratio **(I)**. In tissue samples of 36 HGSOC patients, an optimal cutoff of 3.038 for the S100A8/CRP abundance ratio was found. Patients with the S100A8/CRP abundance ratio higher or lower than the cutoff value were compared. Boxplots show the distribution of the S100A8/CRP abundance ratio after pulling datasets generated in six independent studies **(J)**. See Table S5 for detailed information about patients and samples included in this datasets.. NETs proteins are labeled in orange. Proteins usually linked with local inflammation were marked in blue and in red were labeled protein related to systemic inflammation. PFS - progression-free survival. OSD - overall survival days. * - indicate p-value < 0.05.

In addition, we re-evaluated proteomics data from other laboratories regarding chemotherapy-naive HGSOC patients [48]. Again, NETs proteins were found up-regulated in patients more sensitive to chemotherapy (n = 14, median disease-free survival or PFS = 1160 days) in comparison to chemoresistant patients (n = 11, median PFS = 190 days) (Fig. 5C).

Intriguingly, CRP was not found significantly regulated in both studies. However, the S100A8/CRP abundance ratio was found decreased in the group of patients with poor outcome (in poor responders and patients resistant to chemotherapy, Fig. 5B and 5D). This finding suggested that the validity of S100A8/CRP abundance ratio could be extended as prognostic biomarker from ovarian cancer to melanoma.

However, CT45 (cancer/testis antigen 45) protein was reported to represent the best possible prognostic marker for long-term survival in the ovarian cancer study (Fig. 5 C and D) [48]. Re-grouping the HGSOC patients in this study according to overall survival with OSD < 1000 (n = 10) and OSD > 1000 (n = 15) revealed that S100A8/CRP abundance ratio significantly stratified these groups (Fig. 5F). While CT45 protein seems to better predict chemotherapy response, S100A8/CRP seems to better predict overall survival.

In line, the S100A8/CRP abundance ratio was again found significantly up-regulated in the group of patients with OSD > 1000 as calculated from proteomics data of metastatic tumors isolated from HGSOC patients combining the study of Coscia et al. [48] and a more recent study (Fig. 5G and H, n = 36) [49]. Kaplan-Meier analysis confirmed that patients with an S100A8/CRP abundance ratio equal or above a cutoff value of 3.038 showed again a significantly longer overall survival time compared to patients with values below this cutoff (Fig. 5I). NETs proteins and S100A8/CRP ratio were up-regulated in ovarian cancer patients with favorable outcome of another recent study [50] (Fig. S4A). Thus, we calculated the ratio of the two proteins in a total of six studies (Fig. 1A, 5A–5G, and S4) representing all kinds of variations based on different patient cohorts and methodological details. Indeed, the S100A8/CRP ratio was found significantly correlated with overall survival (n = 116, Fig. 5J, TableS5).

## Discussion

An active and relevant role of neutrophils for tumorigenesis has been recognized with regard to several tumors [51]. However, whether neutrophils rather promote cancer development or contribute to immune reactions inhibiting cancer growth, or whether neutrophils subtypes [52] may account for conflicting data, is still a matter of debate. Here, with regard to HGSOC we have collected ample evidence for NETosis associated with non-miliary tumor spread. NETosis as well as the activity of tumor-associated macrophages have been rather linked to the promotion of metastasis [53, 54]. While non-miliary tumor spread is indeed characterized by more invasive growth, it is associated with better overall survival. Apparently we are dealing here with contradictory observations.

The present data suggest a model which might account for several apparent discrepancies (Fig. 6). The model comprises three levels: 1) initiation of NETosis by hypoxic cell stress; 2) establishment of distinct mascroscopic features related to a specific biomarker profile due to NETosis and 3) modulation of the adaptive immune system by NETosis promoting improved overall survival.

**Figure 6:**
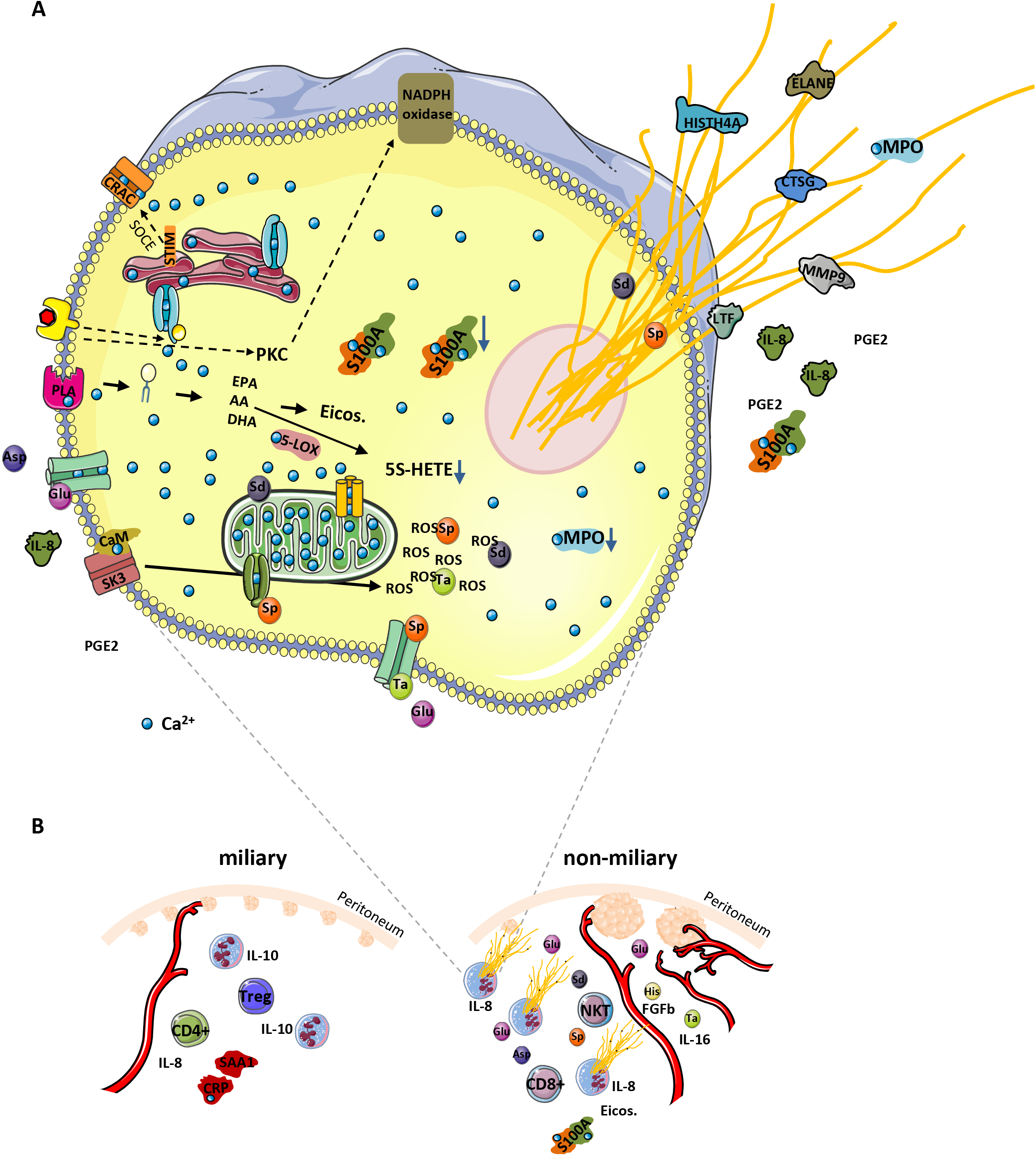
A - Proposed model for NOX-independent NETs formation in HGSOC patients. Increase of cytosolic Ca^2+^ concentration activates and translocates several calcium-dependent and calcium-binding proteins, thus inducing hydrolysis of plasma membrane lipids (releasing PUFAs AA, EPA, and DHA) via activation of a calcium-dependent PLA. Among enzymes (5-LOX, 12-LOX, 15-LOX, COX and CYP450) which metabolize these PUFAs into eicosanoids, only 5-LOX binds Ca^2+^. Under conditions of prolongated high intracellular Ca^2+^ concentration, the activity of 5-LOX enzyme is decreased, resulting in eicosanoid class switching process, exemplified by 5S-HETE. Additionally, elevated Ca^2+^ levels promote translocation of calmodulin to SK3 receptor imbedded into plasma membrane, inducing receptor activation and induction of ROS production from mitochondria resulting in NETosis. The S100A8/9 protein complex is released with the NETs. The sustained Ca^2+^ influx in the cell affects mitochondrial function and may initiate apoptosis. To attenuate this effect, the permeability transition pore channel and Ca^2+^ entry channels, gets closed by spermine (Sp). At the same time, the cell may use all three metabolites (spermine Sp, spermidine Sd and taurine Ta) as ROS scavenger to deal with increased oxidative stress. Both positively charged polyamines stabilize DNA strands, and thus get released together with the NETs. We postulate that glutamine Glu released from cancer cells under hypoxic conditions may induce Ca^2+^ influx in neutrophils by the activation of specific membrane receptors. Glutamine may also promote neutrophil activation by inducing the secretion of IL8 and PGE2. Whereas IL8 is a well-known chemoattractant and inducer of NETosis, PGE2 can induce eicosanoid class switching as observed in ascites samples. The dotted arrows indicate NOX-dependent NETosis and additional pathways via store-operated calcium entry (SOCE) promoting intracellular calcium mobilization. **B – Strong correlation between NETs formation, angiogenesis and the type of tumor spread in HGSOC**, selected proteins, eicosanoids, metabolites, immune cells or processes are depicted apparently regulated in miliary or non-miliray ascites samples. Strongly activated neutrophils, most probably by shedding the millet-like and freshly build small tumor nodules, promote building of bigger but fewer in number of tumor nods in the non-miliary type. In miliary spreading tumors up-regulated IL-10 inhibits NETs formation. Additionally, neutrophils depending on their activation status modulate the immune system by determining the immune cell composition in the tumor microenvironment. Otherwise, increased angiogenesis associated with increased blood supply may contribute to less suppressive effects on neutrophils activation in the non-miliary type. PLA - phospholipase A; Asp – aspartate, His - histamine; calcium release-activated channel (CRAC); PKC - protein kinase C; FGFb - fibroblasts growth factor best; Eicos. - eicosanoids; NTK - natural killer T - cells; Treg - regulatory T cell.

### 1) Initiation of NETosis by hypoxic cell stress

A substantial challenge for ovarian cancer cells in the peritoneum is caused by hypoxia [55] calling for metabolic adaptation [56]. Actually increased incidence of cell death may be associated with stress resulting in the release of DAMPs and the establishment of an adaptive immune response [57]. Also the release of glutamate, presently found up-regulated in non-miliary ascites samples (Fig. 2) has been associated with hypoxic stress of cancer cells [58] potentially causing the initiation of NETosis (Fig. S5) [59]. In addition, the strong chemokine and NETs inducer IL-8 was found up-regulated with glutamate (Fig. S5) and was observed upregulated in non-miliary ascites [11]. Furthermore, IL-8 signaling pathway and IL-8 activated PI3K signaling pathway were identified as one of the most deregulated pathways [60, 61]. As a result, NETosis may account for all major molecular alterations associated with non-milary ascites comprising proteins, eicosanoids and metabolites (Fig. 1 and 6).

### 2) Establishment of distinct mascroscopic features related to a specific biomarker profile due to NETosis

NETs releasing neutrophils eventually release PGE2, IL-8, MMP9, ELANE (Fig. 1A) [62] and other tumor promoters plausibly accounting for a more invasive growth of neighboring tumor cells. An improved access to blood supply resulting from invasion supported by neutrophil-induced neoangiogenesis may account for larger metastases typically observed in case of non-miliary metastases [63, 64]. In contrast to this scenario, ovarian cancer cells better coping with hypoxic stress would rather remain restricted to the peritoneum as observed in case of miliary forms of metastasis, associated with less growth capacity but improved resistance to anti-cancer drugs, less involvement of immune cells and thus decreased overall survival. Additionally, the increased levels of IL-10 (Fig. 2) in miliary samples may account for an inhibition of NETosis [65], thus supporting miliary kind of tumor spread. Indeed, neutrophils are known as main sources of proteins S100 A8 and A9 [31], which may thus serve as biomarkers for NETosis (Fig. 3A).

### 3) Modulation of the adaptive immune system by NETosis promoting improved survival

The involvement of neutrophils in carcinogenesis is well recognized [51], but their role in cancer therapies is still subject of controversial debates [66]. Elevated levels of peripheral neutrophils correlated with poorer clinical outcome in oropharyngeal cancer [67] and have been described to promote metastasis in a mouse lung cancer model [68] as well as human breast cancer [69]. Here, we described that NETs formation is associated with non-miliary metastasis and better overall survival. This may be due to a modulation of the adaptive immune system via recruitment of CD8+ T cells and suppression of regulatory T cells, which were found elevated in miliary samples (Fig. 2). We suggest here that the eicosanoids found deregulated (Fig. 1B, 2) may represent strong effector molecules regulating the local immune status. Most importantly, the present data regarding the prognostic power of the S100A8/CRP ratio seem to suggest a beneficial role of activated neutrophils for cancer patients. This may actually look like data inconsistency or contradicting observations.

A closer look on the present data may resolve this apparent conundrum and put the role of neutrophils for cancer in a new context. It seems to us it is not necessarily the presence or absence of neutrophils which makes the difference. Here we suggest it is the functional state of neutrophils *in situ* which matters. The present data are highly consistent with NOX-independent activation of neutrophils in the peritoneum accompanying non-miliary metastases. The ascites samples actually do not show an up-regulation of LOX-5-derived products, which have been made responsible for tumor-promoting effects of neutrophils [68], but show increased levels of LOX and COX-products associated with the resolution of inflammation. Microparticles containing 14-HDoHE and 17-HDoHE and shedded from neutrophils may inhibit the pro-inflammatory action of activated macrophages [70], which have been demonstrated to promote tumor growth and metastasis [54]. The present *in vitro* studies with neutrophils isolated from healthy donors demonstrate that stimulation via NOX or via calcium release causes substantially different effector functions in neutrophils. While PGE2 released from neutrophils in the periphery may support invasiveness and matastasis [71], local PGE2 release within the peritoneum may stimulate the immune system to better elicit an appropriate immune response. Indeed, increased levels of CD8 positive T-cells were described to be associated with non-miliary metasis (Fig. 2) [11].

In conclusion, both cell activities as well as the localization of specific events seem to matter. Obviously, it is not trivial to seek blood-borne tumor markers reporting specific events confined to specific locations. However, the presently described ratio of S100A8/CRP might provide at least some insight in that regard. Any systemic activation of the immune system including the activation of neutrophils will result in the increase of the highly sensitive and disease-relevant biomarker CRP as demonstrated in countless studies. Increased CRP is typically associated with poor prognosis, in line with all reports regarding the systemic involvement of neutrophils. S100A8, as well as S100A9, are relased by neutrophils upon local NETosis (Fig. 3A). Thus, it is the balance between local inflammation and systemic inflammation which may account for the apparent predictive power of S100A8/CRP ratio for improved survival.

In summary, the present data demonstrate a strong influence of NETosis on the local microenvironment accompanying non-miliary metastasis. The consequences involving the immune system may account for the improved overall survival reported for non-miliary metastases. A new marker profile, the S100A8/CRP ratio was found to correlate with improved survival and may report the individual balance between local and systemic activation of the immune system.

## Materials and methods

Ascites fluid was collected from HGSOC patients at the Medical University of Vienna with ethical approval from The Ethics Committee of the Medical University of Vienna and Vienna General Hospital (AKH), nos. 366/2003 and 793/2011. Metabolomics and Luminex-based cytokinomics data of cell-free ascites, immune cells composition data of ascites and transcriptomics data of tumor tissue were published before [11]. To the 25 cell-free ascites samples (11 miliary, 7 non-miliary and 6 unknown) from the same sample pool, we applied MS-based proteomics and eicosadomics analysis. Additionally, we measured cell-free ascites samples collected from five patients with liver cirrhosis. A co-association network analysis was implemented using all multi-omics data. Further, in-vitro experiments with neutrophils isolated from healthy donors were performed. The experiments with neutrophils were approved by the Ethics Committee of the Medical University of Vienna (number: 1947/2014). Four different software packages were implemented to evaluate the data generated from three mass spectrometric instruments [72, 73] (Fig. S1). An extended description of the material and methods can be found in the supplementary information.

## Acknowledgements

This work was supported by the University of Vienna and by the Joint Metabolome Facility and the Multimodal Imaging Facility of the Faculty of Chemistry of the University of Vienna, members of the Vienna Life Science Instruments (VLSI).

## Author contributions

BM and CG conceived the study. SA, AS and AM provided clinical materials and primary neutrophils. BM, JCM, LM and GDF performed the experiments, BM and CG analyzed the multi-omics data, DP performed the co-association network and Kaplan-Mayer analysis. BM and GC wrote the manuscript which was edited and approved by all the co-authors.

