## Supplemental data for "Neutrophil extracellular trap formation correlates with improved overall survival in ovarian cancer"

### Supplementary data containing

- **Materials and methods**
- **Legends to supplementary figures**
- **Supplementary figures**

#### Materials and methods

*Depletion of ascites samples*

Ascites sample were depleted using Pierce™ Top 12 Abundant Protein Depletion Spin Columns (Thermo Fisher Scientific, USA). The depletion was performed according to the protocol of the manufacturer, using 550 µg total protein amount per depletion.

*Neutrophils isolation and treatment*

Peripheral venous blood was collected using S-Monovete® 7.5 mL K3 EDTA plasma tube (Sarstedt, Germany) from three healthy volunteers. A dextran protocol was implemented for neutrophils isolation [1]. To the 20 mL whole blood was added 10 mL phosphate buffered saline (PBS) and gently rotated. The diluted blood was layered on the top of 15 mL lymphocyte separation medium (PromoCell, Deutschland) and centrifuged under room temperature (RT) condition for 30 minutes (min) at 800g in a HeraeusTM MegafugeTM 16R and BIOShieldTM 720 High-Speed Swinging-Bucket Rotor (Thermo Fisher Scientific). The acceleration was seated to 5 and deceleration was switched off. After aspiration of plasma and peripheral blood mononuclear cells, the pellet of red blood cell with neutrophils on the top was carefully reconstituted in 20 mL PBS. To this mixture was added 20 mL of 6% dextran T500 (Pharmacosmos, Denmark), the falcon tube was gently rotated and incubated for 30 min at RT. The supernatant was collected in a fresh tube and centrifuged at 500g for 5 min in RT, both, acceleration and deceleration were seated at 2. After aspirating the supernatant, three red cell lysis steps were performed and each time the pellet was gently dissolved in 5 mL red cell lysis buffer, incubated for 5 min at RT and after mixing with 20 mL PBS the cell suspension was centrifuged for 5 min at 500g. The composition of red cell lysis puffer: ammonium chloride (154 mM; Merck, Austria), potassium hydrogen carbonate (10mM; Merck, Austria) and EDTA (0.1 mM; ethylenediaminetetraacetic acid disodium salt, Sigma-Aldrich, Austria). In the end, the pellet with isolated neutrophils was washed with 20 mL PBS and centrifuged again for 5 min at 500g. In the very last step the isolated neutrophils were reconstituted in RPMI medium phenol red free (Thermo Fisher Scientific, Austria) supplemented with 10% fetal calf serum (ATCC, USA). After counting the cell were divided into 25cm$^2$ suspension cell culture flasks (Sarstedt, Germany). After 20 min incubation at RT°C and 5% $CO_2$, cells were treated for one hour and three hours with 25 nM phorbol 12-myristate 13-acetate (PMA) and 4 µM Ionomycin (Sigma-Aldrich, Austria). Each time control of untreated cell was prepared. At the end of incubation time, the supernatant was collected with a pipet from the flask and centrifuged at 1100 rpm for 5min. To the cell-free supernatant was added 5 µL of the mixture of eicosanoids deuterated standards and was diluted 1 to 5 with ice cold ethanol for overnight protein precipitation at -20°C. In the next step, the mixture of cell supernatant and ethanol was centrifuged for 40 min at 5000 rpm and 4 °C using a HeraeusTM MegafugeTM 16R and BIOShieldTM 720 High-Speed Swinging-Bucket Rotor (Thermo Fisher Scientific). The pellet containing proteins after drying under vacuum condition was reconstituted in 200µL samples buffer and after the determination of total protein amount, the protein sample of neutrophils supernatant was digested. Sample buffer composition: urea (7.5 M), thiourea (1.5 M), 3-[(3-cholamidopropyl) dimethylammonio]-1-propanesulfonate - CHAPS (4%), sodium dodecyl sulfate - SDS (0.05%) and dithiothreitol - DTT (100 mM). On the other hand, 200 µL of protein-free supernatant were separated for the metabolomics analysis, while the rest of it was used for the analysis of eicosanoids.

For the experiment with glutamate, neutrophils were cultivated in Hank's balanced salt solution (HBSS) medium since the concentration of glutamate in the RPMI medium is very high. The composition of HBSS buffer: $CaCl_2$ (1 mM), $MgCl_2$ x 6H2O (0.5 mM), $MgSO_4$ x $7H_2O$ (4 mM), KCl (5mM), $KH_2PO_4$ (0.4 mM), $NaHCO_3$ (4 mM), NaCl (0.138 M), $Na_2HPO_4$ anhydrous (0.3 mM), D-Glucose (6 mM) and HEPES (0.025 M). For the experiment with glutamate, we followed an experimental design published years ago [2]. The neutrophils were primed with $10^{-11}$ M formyl-met-leu-phe (FMLP) for 15 min. Afterward, cells were treated with 100 µM $Na_3VO_4$ as a phosphatase inhibitor. At the same time to the treated cells was additionally added glutamate (Sigma-Aldrich) to a final concentration of 250 µM, which is the average glutamate concentration determined in non-miliary ascites samples.

*Immunolocalization of spermine/spermidine in neutrophils*

For the subcellular localization of spermine, immunofluorescence experiments were performed. Briefly, after incubation neutrophils were fixed with pre-warmed formaldehyde (FA, 3.7%, 15 min) and permeabilized with Triton X (0.2%, 10 min). Blocking of unspecific binding sites was performed with BSA (Bovine serum albumin 1%, 1h). For the detection of spermine a polyclonal anti spermine antibody was chosen (ab 26975 Abcam, dilution 1:100, 2h), whereas the localization of mitochondria was obtained with an anti TOM20 antibody (Tom20 Antibody F-10 sc-17764, SantaCruz, dil. 1:250). Endoplasmic reticulum was stained with ER staining Kit (ab 139481 Abcam, Green Fluorescence). Primary antibodies were detected with a donkey anti rabbit antibody (Alexa Fluor 568-conjugated A10042 Invitrogen by Thermo Fisher Scientific, dil. 1:1000, depicted in red) and a donkey anti mouse (Alexa Fluor 647-conjugated A31571 Invitrogen by Thermo Fisher Scientific, dil. 1:1000, depicted in white), respectively. After multiple washing steps, neutrophils were post-fixed with FA 3.7% (10 min) and reactive sites were masked with glycine (100 mM). Slides were mounted with Roti-Mount FluoCare with DAPI to counterstain the nuclei (Roth, Graz, Austria). Images were acquired with a Confocal LSM Zeiss 710 equipped with ELYRA PS. 1 system with a Plan Apochromat 63X/1.4 oil objective.

*Isolation of mitochondria and endoplasmic reticulum from neutrophils*

Endoplasmic reticulum (ER) and mitochondria were isolated using differential sucrose gradient [3]. Neutrophils from a healthy donor were isolated as described above and treated for one hour with PMA and ionomycin. Cells were scratched directly in the flask without removing the medium, transformed into a fresh tube, and pelleted under could condition (4°C) by centrifugation at 1100 rpm. The cell pellet was resuspended in 1 mL lysis buffer. The lysis buffer consists of: D-mannitol (220 mM), sucrose (60 mM), HEPES (20 mM), $MgCl_2$ (1.5 mM), and EDTA (1 mM). After one freeze and thaw cycle in liquid nitrogen, the cells were lysed using a 20G needle and followed by a centrifugation step at 700g to remove the large cellular debris. The supernatant from the last centrifugation step was again centrifuged at 15 000g and 4°C to separate crude mitochondria (pellet) from crude ER. Ultracentrifugation was performed using an Avanti$^{TM}$ centrifuge J-30 I (Beckman coulter, USA), a JS-24 rotor (Beckman, USA) and ultra-clear centrifugation tubes (16 x 96 mm) (Beckman, USA). The sucrose gradient for ER isolation consists of 2 mL 2M, 3 mL 1.5M and 3 mL 1.3M, whereas for mitochondria isolation two stage gradient consisting of 1 mL 1.7M and 1.6 mL 1M was used. The supernatant containing the crude ER was carefully pipetted on the top of the sucrose gradient and centrifuged for 100 min at 110 000g and 4°C. In the interface of the 1.3 M of sucrose gradient layer using a needle and syringe were removed 0.6 mL volume, which was collected in a fresh centrifugation tube, diluted with 3.8 mL lysis buffer and centrifuged for

45 min at 110 000g. The supernatant was discard and the tube was inverted to dry the pellet with isolated ER, which was finally reconstituted in 50 µL PBS and stored at -20°C. The pellet with crude mitochondria was resuspended in 0.6 mL lysis buffer and was layered in the top of the sucrose gradient. The mitochondrial gradient was centrifuged for 22 min at 40 000g and 4°C. Afterward, in the interface between 1.7 M and 1 M of sucrose layers was removed 0.4 mL volume using a syringe and after dilution with 1.5 mL lysis buffer was centrifuged for 10 min at 15 000g. The supernatant was discarded and the pellet with isolated mitochondria after drying was reconstituted in 50 µL PBS and stored at -20°C. The entire volume of ER and mitochondria samples were used for the metabolomics analysis of spermine and spermidine as described below.

*In-solution digestion*

In-solution digestion was performed using 10 kDa MW cut-off filters (Nanosep with Omega membrane, Pall, USA). For both samples, depleted ascites samples and supernatant of neutrophils, 20 µg total protein amount were used for digestion. The digestion protocol consists of protein reduction with 32mM dithiothreitol (Gerbu), protein alkylation with 54 mM 2-iodoacetamide solution (Sigma Aldrich) and overnight tryptic digestion, as described recently [4]. A Trypsin/Lys-C Mix (MS grade; Promega) was used for digestion with a total enzyme to protein ratio of 1:20. For clean-up of resulting peptide samples C18 spin columns (Pierce™ C18 Spin Columns, Thermo Fisher Scientific, USA) were used and, after drying via vacuum centrifugation, the samples were stored at -20°C. Upon LC-MS analysis the dried peptides were reconstituted in 5 µL of the equimolar 10 fmol standard peptide mix and 40 µL of mobile phase A (98% H2O, 2% ACN, 0.1% FA) for shotgun analysis, and for targeted analysis in 30 µL of the 10 fmol/µL standard peptide mix solution containing 30% formic acid.

*Untargeted shotgun LC-MS proteomics analysis*

Shotgun proteomics analysis of digested peptide was conducted on a QExactive Orbitrap mass spectrometer (Thermo Fischer Scientific) coupled with a UltiMate 3000 RSLC nano System (Pre-column: Acclaim PepMap 100, C18 100 µm x 2 cm; Analytical column: Acclaim PepMap RSLC C18 75 µm x 50 cm; Dionex, California) [4]. A 54 min gradient from 1% to 40% solvent B was applied for chromatographic peptide separation. Solvent composition: A – 97.9% water, 2% acetonitrile, 0.1% formic acid and B – 97.9% acetonitrile, 2% water and 0.1% formic acid. The resolution on Orbitrap mass spectrometer was set to 70,000 for full MS scan and the MS2 scan was acquired at a resolution of 17,500, both at m/z 200. The MaxQuant 1.5.2.8 [5] software packet was used for protein identification and MS1 based label-free protein quantification (LFQ) (Table S1). The same criteria as described recently were applied for protein identification and MS1 based protein qualification [4]. Briefly, search criteria included a maximum of two missed cleavages and a maximal mass deviation of 5 ppm for peptide ions and of 20 ppm for fragment ions. Further, a minimum of two peptide identifications per protein (including one unique) was requested and an FDR of less than 0.01 was applied at both peptide and protein level. For statistical analysis, data obtained from both biological as well as technical replicates were included and the Perseus statistical analysis package was used[6] (Table S1).

*Targeted LC-MRM-proteomics analysis*

Proteomics targeted multiple reaction monitoring (MRM) analysis was performed on an Agilent 6490 triple quadrupole mass spectrometer coupled with a nano-Chip-LC Agilent Infinity Series HPLC1260 system, as described recently [4, 7]. Solvent compositions were 97.8% H2O, 2% ACN and 0.2% FA for solvent A, and 97.8% ACN, 2% H2O and 0.2% FA for solvent B. We developed an MRM method for the measurement of selected proteins by applying an established workflow [4, 7](Table S3). The skyline software (v. 4.1.0.11796) was used for method development and data evaluation [8]. Data were normalized to four synthetic standard peptides.

*Bone marrow plasma samples collected from multiple myeloma patients*

Plasma samples were obtained from routinely taken bone marrow aspirates at the Vienna General Hospital in course of a published study [9]. Written informed consent was obtained from all patients, as well as approval of the Ethics Committee of the Medical University of Vienna (application

nr. 1181/2013). All samples preparation steps and the parameters of the mass spectrometry-based proteomics analysis were similar to those applied for the analysis of ascites samples.

*Eicosanoid extraction*

Proteins were precipitated overnight with ethanol from 200 µL ascites sample after adding of deuterated internal standard (15S-HETE-d8 and PGF2a-d4). Samples were than centrifugation for 40 min at 5000 rpm and 4 °C using a HeraeusTM MegafugeTM 16R and BIOShieldTM 720 High-Speed Swinging-Bucket Rotor (Thermo Fisher Scientific). The collected supernatants were preceded for eicosanoid extraction using StrataX solid-phase extraction (SPE) columns (StrataTM-X 33 µm polymeric reversed phase, sorbent mass 30 mg/1 mL, phenomenex®). The SPE column was first washed with 2mL methanol (VWR International) and conditioned with 2 mL water (VWR International). The sample was loaded to an SPE column after diluting the organic phase to 10% with water. A wash step with 2 mL water was performed before eluting the eicosanoids from the SPE column with 500 µL methanol supplemented with 2% formic acid. The eicosanoid extract was dried completely in a SpeedVac (Genecav™) at 35°C. Afterward, samples were reconstituted in 200 µL LC-MS starting conditions (35 % B, 65 % A) and immediately analyzed via LC-MS.

Stock solutions of deuterated standards had a concentration of 25 µg/mL. A deuterated standard mix of 4 µL of 15S-HETE-d8, 12S-HETE-d8, 11,12-DiHETrE-d11 and 20-HETE-d6 as well as 12 µL of 5-Oxo-ETE-d7 and 8 µL of PGE2-d4 was diluted to 500 µL with ACN. Out of this standard mix, 5 µL was added to each sample. The standard mix of deuterated eicosanoids was added to the supernatant samples upon harvesting of 3 mL supernatant from cultured neutrophils isolated from healthy donors, followed by overnight protein precipitation with ethanol at -20°C. By centrifugation for 40 min at 5000 rpm and at 4 °C using a HeraeusTM MegafugeTM 16R and BIOShieldTM 720 High-Speed Swinging-Bucket Rotor (Thermo Fisher Scientific), the supernatant was separated from the protein pellet. The volume of supernatant sample was reduced to 1.5 mL in a SpeedVac (Genecav™) at 35°C and diluted with water to 15 mL. The same protocol was followed for the SPE eicosanoid extraction as described above. In the end, the samples were reconstituted in 150 µL LC-MS starting conditions (35 % B, 65 % A) and immediately analyzed via LC-MS.

*Untargeted eicosadomics LC-MS analysis*

The LC-MS analysis of eicosanoids extracted from ascites samples was conducted in a Q-Exactive Orbitrap mass spectrometer (Thermo Fisher Scientific) using the HESI-source to achieve negative ion mode ionization and coupled with an Infinity 1290 UHPLC system from Agilent, as described previously [4, 10, 11]. Each time 20 µL sample were injected to a Kinetex® C18 column (Kinetex® 2.6 µm C18 100 Å, LC Column 150 x 2.1 mm (phenomenex®). Eicosanoids were separated by applying a 35 min gradient from 35% B to 95% B to a UHPLC system operating at 450 µL/min flow rate. Raw files were analyzed manually using Thermo Xcalibur 2.2 Sp1.48 (Qual browser). Libraries from Lipid Maps depository were used and implemented as references for MS/MS-based identification of eicosanoids. Further, synthetic standards were used for verification of identified eicosanoids. The skyline software was applied for the abundance assessment of identified eicosanoids at MS1 level and the data were normalized to deuterated standards, PGF2-alpha-d4 and 15S-HETE-d8 (Table S2).

*Targeted LC-MRM-eicosadomics analysis*

An Agilent 6490 triple quadrupole mass spectrometer coupled with an HPLC Agilent Infinity Series HPLC1290 system was used for the targeted MRM-eicosadomics analysis. A Kinetex® C18 column (Kinetex® 2.6 µm C18 100 Å, LC Column 150 x 2.1 mm (phenomenex®) was used for the separation of eicosanoids extracted from the supernatant of healthy neutrophils by applying a 10 min gradient from 35% B to 90% B and flow rate of 250 µL/minute. Eluent A consisted of H2O + 0.2 % FA, eluent B of 10 % MeOH + 90 % ACN + 0.2 % FA. Each time 20 µL sample were injected into the HPLC system. Synthetic eicosanoid standards were used for the MRM-eicosadomics assay development [10] (Table S4). Skyline software was implemented for method development and data evaluation [8]. The data were normalized to the global standard, except for PGE2, 20-HETE, 15S-HETE, 12S-HETE and 5-oxo-ETE, which were normalized to respective deuterated standards. Total area normalized (TAN) represent the sum of peak area overall transitions for an eicosanoid normalized to the sum of the peak area of all measured transitions of either global standards or only respective deuterated standard.

*Targeted metabolomics analysis*

Targeted metabolomics of ascites samples was performed using the AbsoluteIDQ p180 kit (Biocrates Life Sciences AG, Innsbruck, Austria) [12]. The kit allows the identification and (semi-) quantitation of metabolites including acylcarnitines, amino acids and biogenic amines, the sum of hexoses, sphingolipids and glycerophospholipids by LC-and flow injection analysis (FIA)-MRM. The samples were analyzed on an AB SCIEX QTrap 4000 mass spectrometer using an Agilent 1200 RR HPLC system, which were operated with Analyst 1.6.2 (AB SCIEX). All amino acids and biogenic amines were derivatized with phenylisothiocyanate. The experiments were validated with the supplied software (MetIDQ, Version 5-4-8-DB100-Boron-2607, Biocrates Life Sciences, Innsbruck, Austria). In the supernatant samples of healthy neutrophils were measured only six metabolites (spermine, spermidine, taurine, histamine, aspartate and glutamate), which build a hub in the middle of the network signature, by implementing the protocol of Biocrates kit for sample preparation as well as for the LC-MS analysis. Proteins from the cell supernatant were precipitated overnight with ethanol and after centrifugation 50 µL of remaining supernatant were used for metabolomics analysis. The skyline software was used for the evaluation of metabolomics data generated from the cell supernatant samples [8]. The data were normalized to the respective deuterated standards, except for histidine, where the deuterated taurine standard was used for data normalization.

*Co-association network analysis*

For the integration of the eicosanoid results with other omics and medium-dimensional data from high grade serous ovarian cancer patients following previously published results of (partly) the same patients were used: metabolomics data of cell-free ascites (AbsoluteIDQ p180 kits, Biocrates)[13], cyto- and chemokines of cell-free ascites (Luminex)[12, 14], immune-cell populations in ascites, serum, and tumor tissues (FACS and immunofluorescence data of ascites immune cells)[12]. The datasets were normalized and log2 transformed as described in the corresponding papers. For each dataset and the eicosanoids the significantly deregulated analytes (FDR < 5%) between samples of patients with miliary and non-miliary peritoneal tumor spread were determined with linear modelling using the R-package limma 3.40.2 [15] and used to label analytes in the network shown in Figure 2 as larger sized nodes. Starting with the list of analysed eicosanoids for each eicosanoid the significantly correlated analytes from all other datasets were determined by linear modelling with limma using the eicosanoid concentrations as independent variables. The significance cutoff for these correlations was set to FDR < 20%. All significant correlations between two analytes if at least one of the two connected analytes was significantly deregulated between miliary and non-miliary were collected and combined to a final correlation network, indicating positive correlations with red and negative correlations with blue arrows. Node colors represent the $\log_2$ fold-change between miliary (up=red) and non-miliary (up=green) samples. The final network was rendered by a force-directed graph drawing methode following Kamada and Kawai [16] implemented into the *network* 1.15 R-package [17].

*Kaplan-Meier analysis*

The optimal cutoff for the Kaplan-Meier estimate shown in Figure 5I was calculated with the cutp-function [18] from R-package survMisc 0.5.5 and the p-value calculated with Cox regression.

All calculations and analyses were performed in R 3.6.0 (R Core Team (2019) [19].

**Figure Legends – supplemental data**

**Figure S1: Workflow of multi-omics analysis.** The multi-omics analysis was performed with ascites samples and supernatant of neutrophils. In each sample proteins, eicosanoids and metabolites were measured. Shotgun proteomics analysis of depleted ascites samples following protein digestion was conducted on a Q Exactive Orbitrap mass spectrometer coupled with a nano-LC system. MaxQuant software tools were used for protein identification and MS1 based protein quantification. After precipitation of proteins with ethanol from ascites samples, eicosanoids were isolated using SPE extraction technique. Eicosadomics screening analysis was performed on a Q Exactive Orbitrap mass spectrometer coupled with a UHPLC system. Eicosanoids were manually identified based on MS/MS spectra using Xcalibur software. However, the correct eicosanoid identification was confirmed with synthetic standards. Skyline software was used for abundance assessment at MS1 level. The Biocrates kit was implemented for the analysis of metabolites. A targeted multi-omics approach conducted on triple quadrupole mass spectrometer was applied for the analysis of supernatant samples isolated from neutrophils. All targeted multi-omics data were evaluated with Skyline software.

**Figure S2: Results of eicosadomics analysis with ascites samples.** Boxplots represent the abundance of selected eicosanoids in ascites samples. In the y-axis in logarithmic scale is plotted the abundance as peak area normalized to global standards.

**Figure S3: Results of ecosadomics analysis with supernatant samples of neutrophils isolated from healthy donors.** Neutrophils were cultured in a medium supplemented with 10% FCS. Cells isolated from healthy donors (n=3) were treated with PMA (25nM) or ionomycin (4µM) and a targeted MS approach was implemented for the analysis of the supernatant samples **(A)**. Neutrophils (n = 5 ) were treated only with 4µM ionomycin for 15 minutes, 1 hour or 3 hours. The data were generated using a Q Exactive HF mass spectrometer by applying an untargeted approach, and the abundance was assessed based on MS1 spectra **(B)**. Error bars indicate standard deviation.

**Figure S4: Shotgun proteomics.** Data were generated by the analysis of tissue samples taken from ovarian cancer patients **(A)** and bone marrow plasma samples of multiple myeloma patients **(B)**. The volcano plots represent protein differences between compared patients, while boxplots show the distribution of the ratio. A more favorable outcome characterizes endometrioid carcinoma (EC) compared to HGSOC. MGUS is a pre-stage of multiple myeloma (MM). * - indicate p-value < 0.05.

**Figure S5: Targeted multi-omics analysis of neutrophils treated with glutamate.** Neutrophils isolated from a healthy donor were first primed by 15 minutes treatment with $10^{-11}$ M FLMP. After that, the control cells were treated only with 100µM $Na_3PO_4$ for 3 hours and to the other cells were additionally added Glu to the final concentration of 250µM. The TAN values for both applied treatment conditions of molecules of interest were shown **(A)**. The fold changes for Spe and Spd in Glu treated compared to control samples **(B)**.

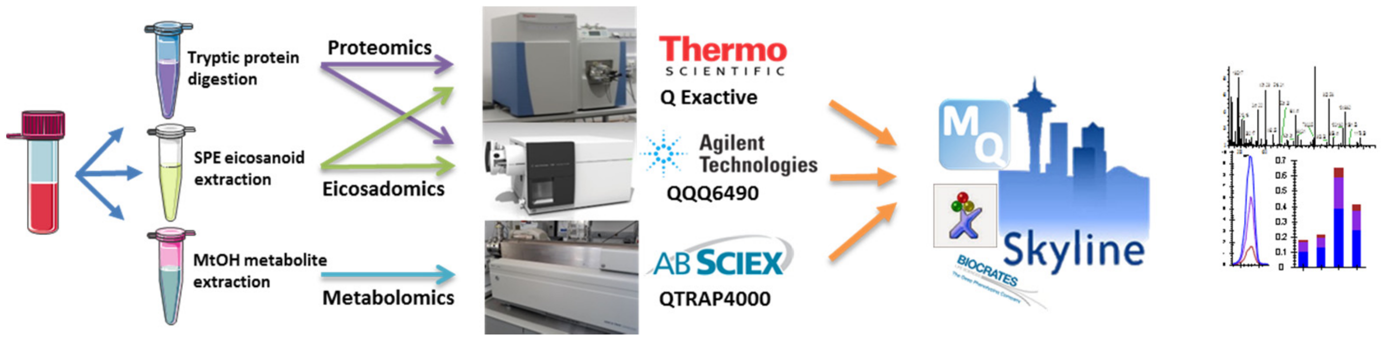
Tryptic protein digestion
SPE eicosanoid extraction
MtOH metabolite extraction
Proteomics
Eicosadomics
Metabolomics
Thermo SCIENTIFIC
Q Exactive
Agilent Technologies
QQQ6490
AB SCIEX
QTRAP4000
MQ
Skyline
BIOCRATES

**Figure S1.**

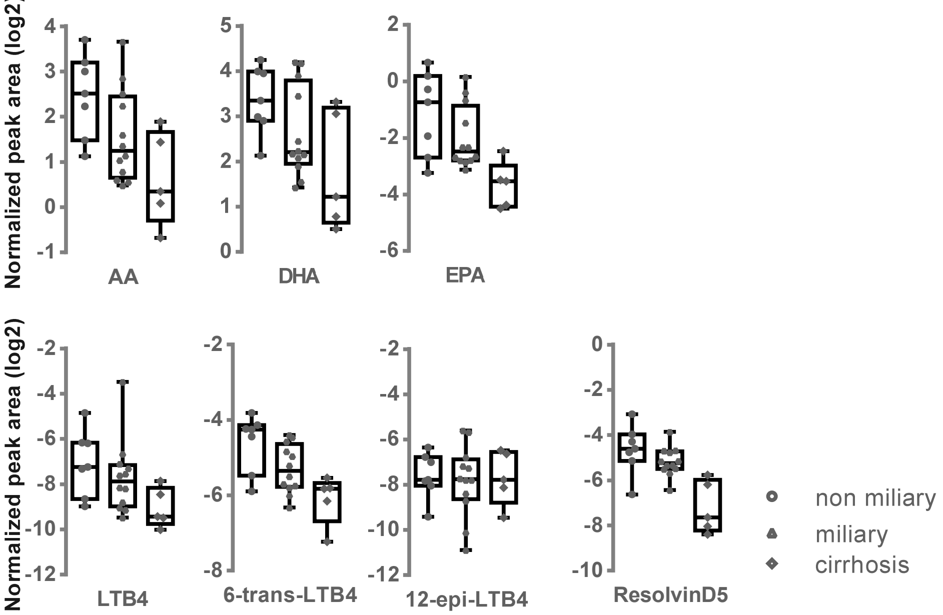
Normalized peak area (log2)
AA
DHA
EPA
LTB4
6-trans-LTB4
12-epi-LTB4
ResolvinD5
non miliary
miliary
cirrhosis

**Figure S2.**

A

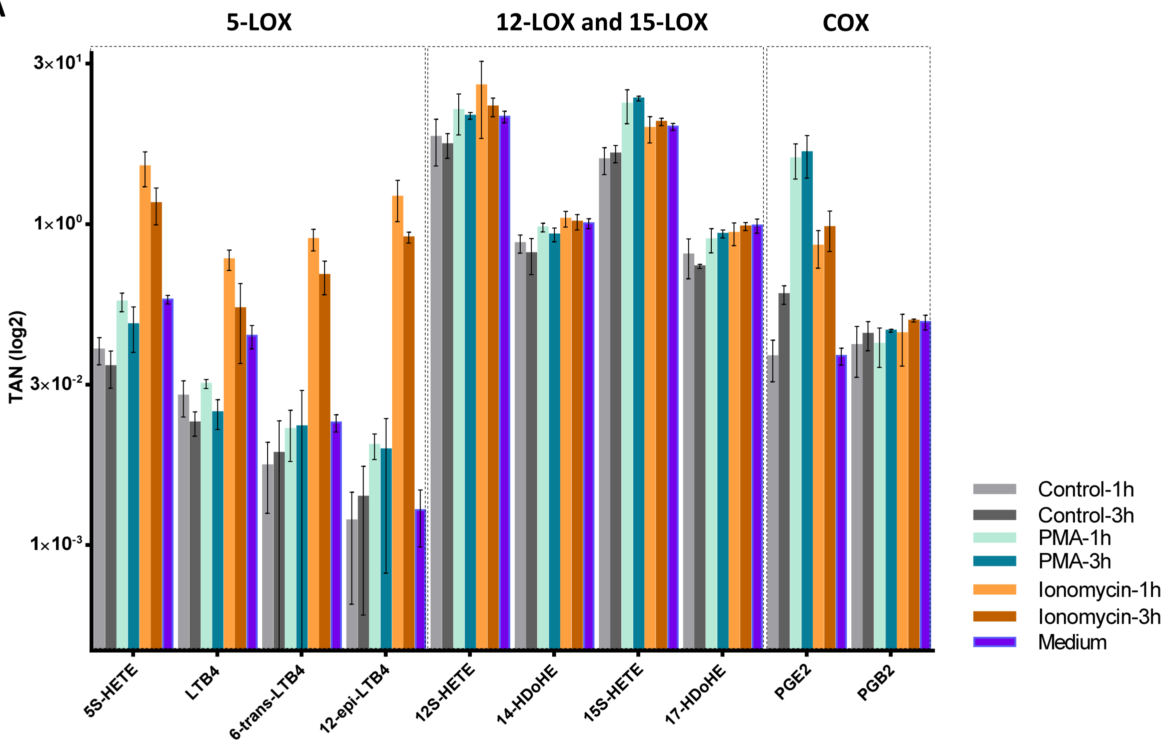
5-LOX
12-LOX and 15-LOX
COX
3×10¹
1×10⁰
3×10⁻²
1×10⁻³
TAN (log2)
5S-HETE
LTB4
6-trans-LTB4
12-epi-LTB4
12S-HETE
14-HDoHE
15S-HETE
17-HDoHE
PGE2
PGB2
Control-1h
Control-3h
PMA-1h
PMA-3h
Ionomycin-1h
Ionomycin-3h
Medium

B

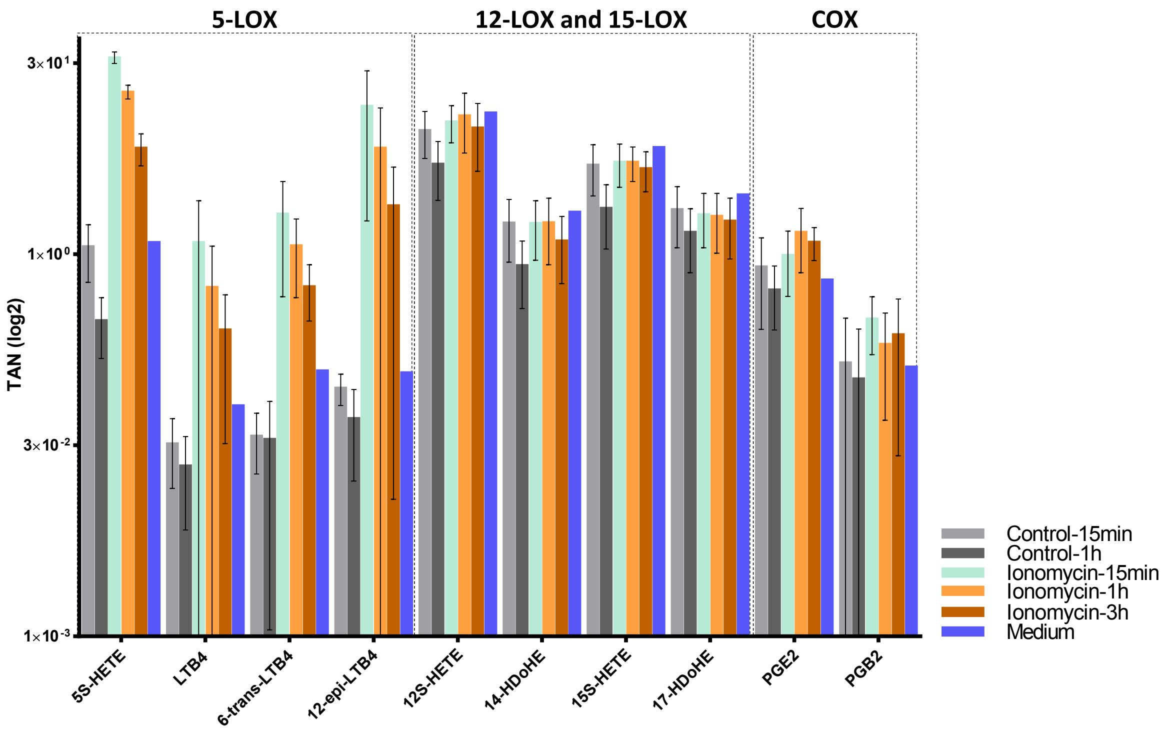
5-LOX
12-LOX and 15-LOX
COX
3×10¹
1×10⁰
3×10⁻²
1×10⁻³
TAN (log2)
5S-HETE
LTB4
6-trans-LTB4
12-epi-LTB4
12S-HETE
14-HDoHE
15S-HETE
17-HDoHE
PGE2
PGB2
Control-15min
Control-1h
Ionomycin-15min
Ionomycin-1h
Ionomycin-3h
Medium

**Figure S3.**

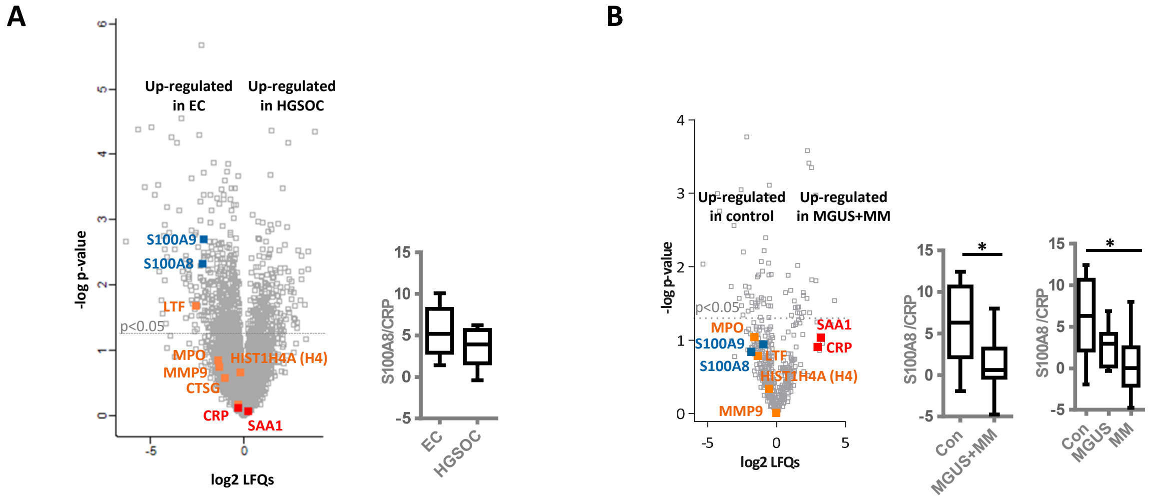
A
Up-regulated in EC
Up-regulated in HGSOC
-log p-value
log2 LFQs
p<0.05
S100A9
S100A8
LTF
MPO
MMP9
CTSG
HIST1H4A (H4)
CRP
SAA1
S100A8/CRP
EC
HGSOC
B
Up-regulated in control
Up-regulated in MGUS+MM
-log p-value
log2 LFQs
p<0.05
SAA1
CRP
MPO
S100A9
S100A8
LTF
HIST1H4A (H4)
MMP9
S100A8 /CRP
Con
MGUS+MM
*
S100A8 /CRP
Con
MGUS
MM
*

**Figure S4.**

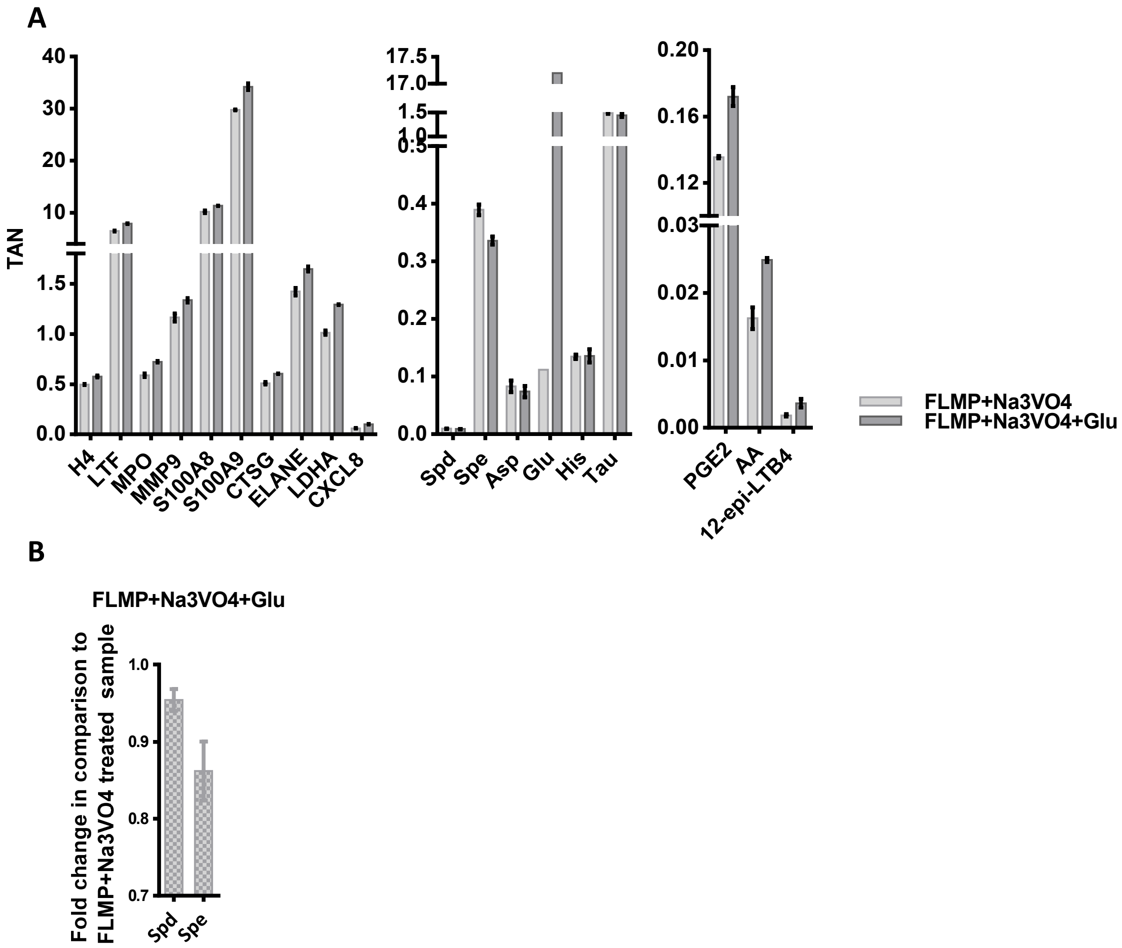
A
TAN
H4
LTF
MPO
MMP9
S100A8
S100A9
CTSG
ELANE
LDHA
CXCL8
Spd
Spe
Asp
Glu
His
Tau
PGE2
AA
12-epi-LTB4
FLMP+Na3VO4
FLMP+Na3VO4+Glu
B
FLMP+Na3VO4+Glu
Fold change in comparison to FLMP+Na3VO4 treated sample
Spd
Spe

**Figure S5.**
